# Host-microbiome transplants of the schistosome snail host *Biomphalaria glabrata* reflect species-specific associations

**DOI:** 10.1101/2023.02.01.526614

**Authors:** Ruben Schols, Isabel Vanoverberghe, Tine Huyse, Ellen Decaestecker

**Author notes:** Corresponding author &.

## Abstract

Snail-borne diseases affect more than a quarter of a billion people worldwide and pose a high burden in the livestock industry. A fundamental understanding of the drivers of the epidemiology of these diseases is crucial for the development of sustainable control measures. The microbiome is increasingly being recognized as an important player in the tripartite interaction between parasitic flatworms, snail intermediate hosts and the snail microbiome. In order to better understand these interactions, transplant experiments are needed, which rely on the development of a reliable and reproducible protocol to obtain microbiome-disturbed snails. Here we report on the first successful snail microbiome transplants, which indicate that *Biomphalaria glabrata* can accrue novel bacterial assemblies depending on the available environmental bacteria obtained from donor snails. Moreover, the phylogenetic relatedness to the donor significantly affected the survival probability of the recipients, corroborating the phylosymbiosis pattern in freshwater snails. The transplant technique described here, complemented by field-based studies, could facilitate future research endeavors to investigate the role of specific bacteria or bacterial communities in parasitic flatworm resistance of *B. glabrata* and might ultimately pave the way for microbiome-mediated control of snail-borne diseases.

## Main text

Evidence is mounting that environmental factors affect the community of host-associated microorganisms and their gene functions (microbiomes). In turn, they can play an important role in ecosystem functioning by modifying host phenotypes [1–4]. The microbiome is known to affect the host’s physiology, behavior and disease resistance [5]. The latter is pertinent for the transmission potential of diseases [6] and changes in the microbiome of intermediate hosts may thus indirectly affect ecosystem health [7]. Intermediate hosts, parasites and microbes interact to determine infection outcome and, subsequently, the potential for disease transmission to definitive hosts [8]. There are precedents with mosquito-borne diseases, such as malaria, dengue and chikungunya, where the release of microbiome-altered vectors reduced disease prevalence, both at a local and at a regional scale [3, 9, 10]. It is hypothesized that microbiomes might affect the outcome of parasitic flatworm infections in snails by competing for resources, producing antimicrobials or stimulating the host’s immune response [11–14].

Such findings have been stimulating schistosomiasis research to elucidate the tripartite interaction between the freshwater snail host, its microbiome and its parasitic flatworm community. Schistosomiasis is a parasitic disease caused by schistosome flatworms (Digenea, Platyhelminthes). It burdens millions of people and animals globally due to inefficient control measures [15]. Different snail-parasite strain combinations exhibit various reciprocity to infection, of which, the underlying pattern is often ill-understood [13, 16]. One intermediate host, the Brazilian snail *Biomphalaria glabrata* is a model system for the study of schistosomiasis. Consequentially, it became the target for research focused on the snail-microbiome-flatworm interaction [11–13]. A decades-long gap exists with earlier publications describing and manipulating the bacteria of *B. glabrata* [17, 18]. These works, however, provide the foundation for current efforts on host-microbiome interactions, more in particular for microbiome transplants.

Transplant experiments are manipulations whereby a number of bacterial strains, or entire bacterial communities are transplanted from a host or substrate to another host [5]. These experiments help us to decipher the host-bacteria interaction and showed, for example, that the host-associated bacterial community determines bumblebee resistance to parasites [1], water flea tolerance to toxic cyanobacteria [2] and mice’s tendencies to become obese [4]. Such transplant experiments, however, depend on the development of a reliable and reproducible protocol to obtain microbiome-disturbed recipients. The current study describes the optimization of the protocol designed by Chernin in 1957 [17], its thorough validation with molecular tools, and the first successful transplants of bacterial communities across a taxonomic range of freshwater snails.

The experimental design aimed to test 1) whether microbiome-disturbed snails would acquire a donor microbiome, 2) what the optimal timing for exposure is, and 3) how the donor-recipient phylogenetic relationship affects the transplant outcome (**Fig. 1**). Snails were bleach-washed at different timepoints (three days, two days and one day prior to, and on the same day as, receiving a donor inoculum; Supplementary Information: “Transplant experiment”, **Fig. S1**). Microbiome-disturbed *B. glabrata* individuals were exposed to three donor inocula isolated from *B. glabrata* individuals (different maternal line), *Planorbarius corneus* individuals, and *Lymnaea stagnalis* individuals. Bleach-washed individuals were tested for the presence of bacteria through a growth assay, microscopic investigation and qPCR (Supplementary Information: “Validating axenicity”). The bacterial communities were characterized through 16S metabarcoding (Supplementary Information: “16S metabarcoding”). Cell- and DNA-based mock communities were included to minimize pipeline biases (Supplementary Information: “Mock communities”; **Fig. S6, Table S2**) [19]. The data processing pipeline is outlined in Supplementary Information: “Data analyses”.

**Figure 1:**
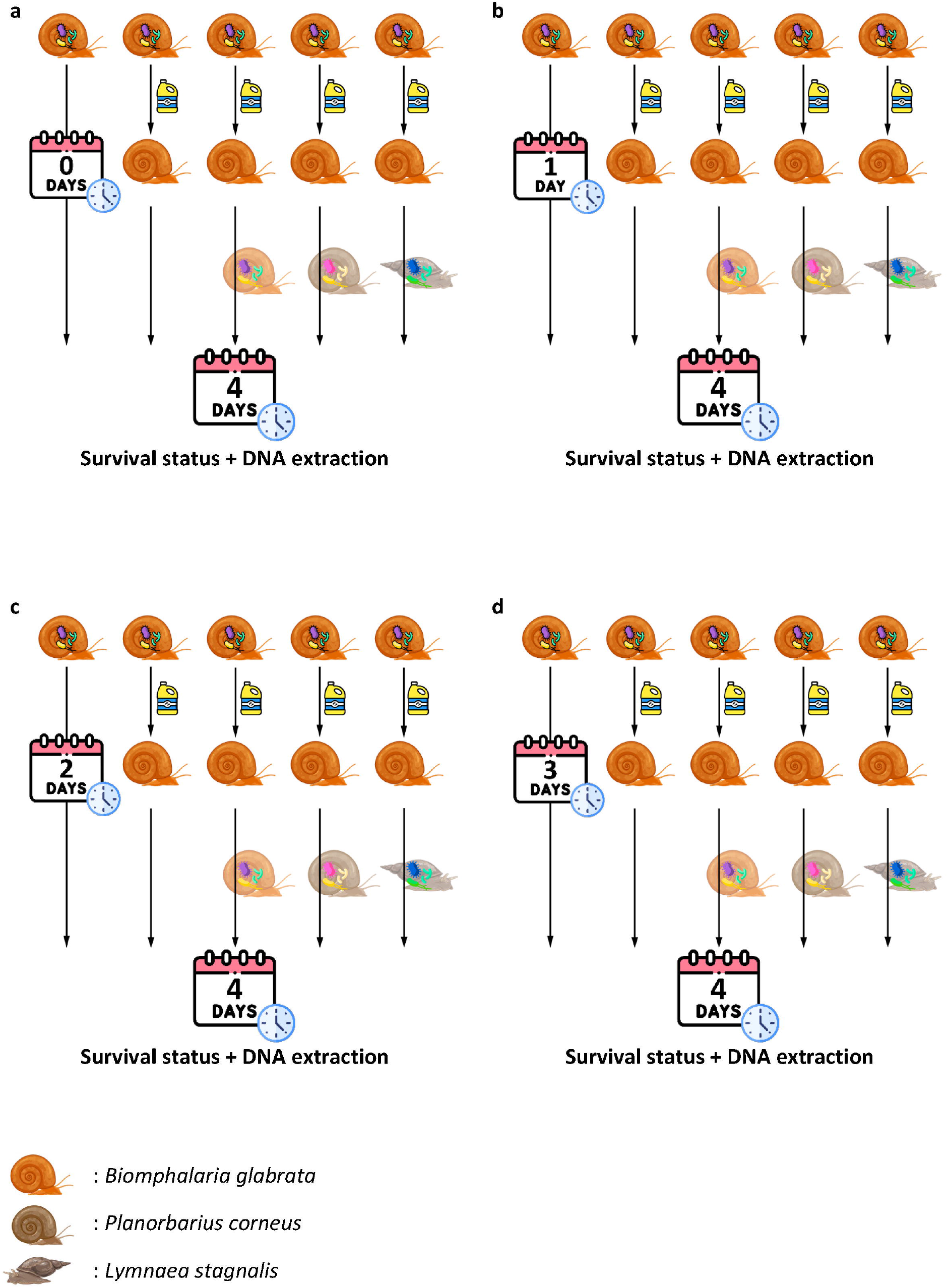
Experimental setup. Snails were sterilized on (**a**) the same day as, and (**b**) one day, (**c**) two days and (**d**) three days prior to, receiving a donor inoculum (see Supplementary Information for detailed manipulations). Microbiome-disturbed *Biomphalaria glabrata* individuals (n=8) of three maternal lines (n=3) were exposed to three donor inocula isolated from: *B. glabrata* individuals (same species but different maternal line), *Planorbarius corneus* individuals (Family: Planorbidae), and *Lymnaea stagnalis* individuals (Superorder: Hygrophila). Additionally, for each factorial combination of time (n=4) and maternal line (n=3), positive (dissected without bleach exposure) and negative (dissected with bleach exposure but no donor inoculum) controls were included (four technical replicates). Survival status of the egg/juvenile was noted on the day of the donor inoculum exposure, three days post donor inoculum exposure and four days post donor inoculum exposure when the specimens were sacrificed for DNA extraction.

Axenicity was confirmed for five out of a total of twelve samples (**Fig. S2, Table S1**) and was affected by the maternal line of the recipient (**Fig. S3**, p<0.001). The donor inoculum affected overall survival chance (**Fig S4**, p<0.001). Moreover, snails that received a *P. corneus* donor inoculum survived better than snails without inoculum or with a *L. stagnalis* donor inoculum (p=0.02). A similar trend is notable for snails that received a *B. glabrata* donor microbiome (p=0.07). The combination of all these results corroborates the phylosymbiosis pattern reported by Huot et al. [11]. Sterilization at the same day as exposure to a donor inoculum induced a high mortality rate (**Fig. S5**).

Microbiome-disturbed samples, negative controls and the *L. stagnalis* donor inoculum had a lower alpha diversity compared to the other samples (**Fig. S7**, p<0.01). The type of donor inoculum affected the bacterial community composition of the recipients (**Fig. 2** and **Fig. S8**, p<0.005). **Fig. 2a** shows each treatment type to cluster most closely to its donor microbiome yet more distant from the centroid. This can potentially be explained by 1) the ‘Anna Karenina principle’, whereby dysbiotic communities tend to be more dissimilar than healthy communities or 2) alternative stable states of bacterial communities [20]. Our data seems to indicate the latter hypothesis as most probable, because individuals that received the same bacterial inoculum display lower dissimilarities compared to controls (**Fig. 2b, Fig. S9**) [20]. To discern between both hypotheses future experiments should assess the stability and variability of the bacterial community, following a transplant event, after a small perturbation and over time [20]. The latter should also reveal how long the bacterial transplants persist over time.

**Figure 2:**
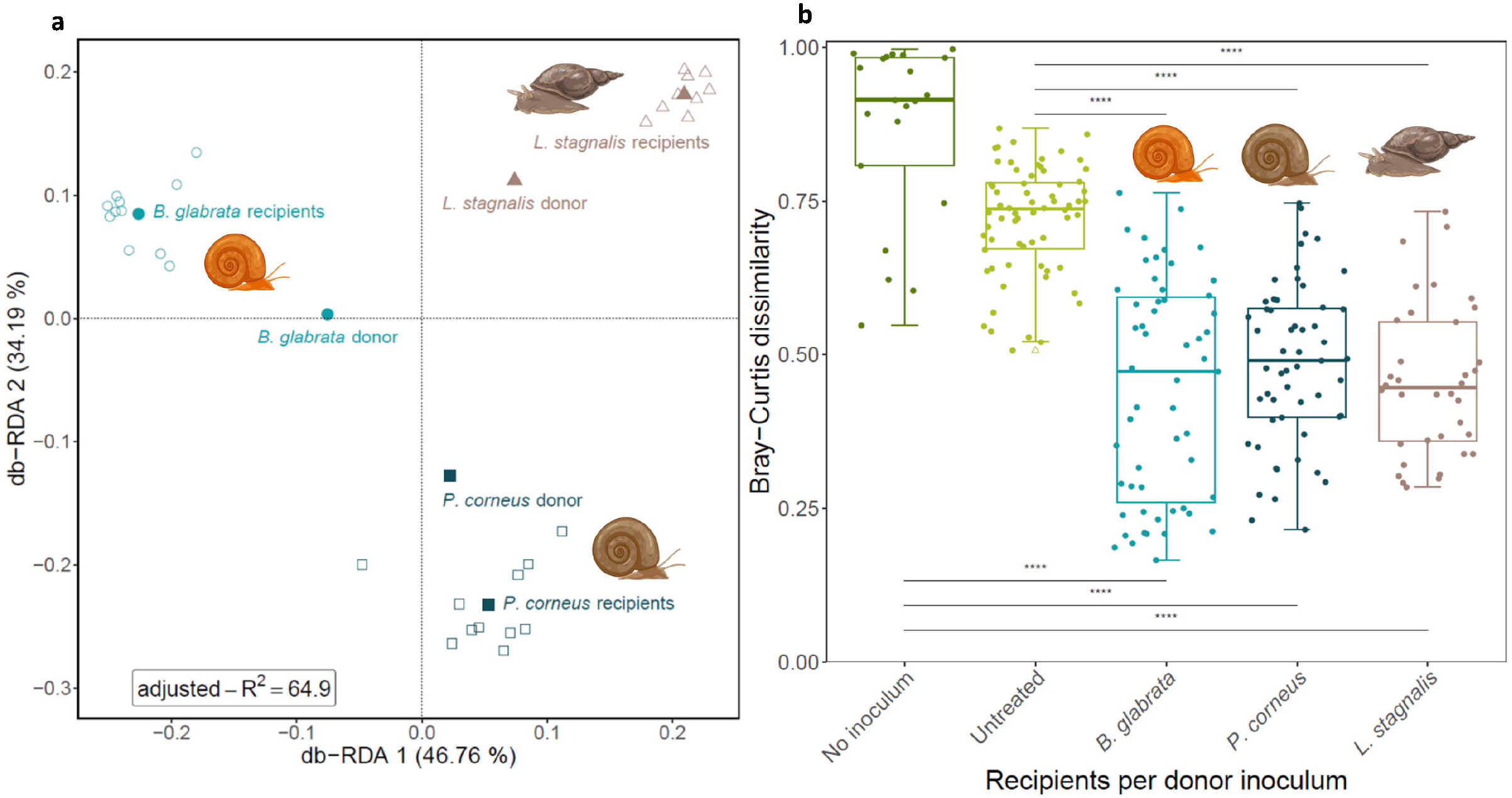
Beta diversity measures across the different treatments. **a** RDA plot showing the beta diversity of the samples that received a donor inoculum (Bray-Curtis dissimilarity). The model contained only the donor type and had an R^2^ value of 64.9%. **b** The within-group Bray-Curtis dissimilarity (n=233). The abbreviations refer to the following: GLA (*Biomphalaria glabrata* recipients), LYM (*Lymnaea stagnalis* recipients), neg (microbiome-disturbed specimens), PLA (*Planorbarius corneus* recipients) and pos (untreated specimens). Outliers are indicated by triangles. The lower and upper hinges correspond to the first and third quartiles (the 25th and 75th percentiles). The whiskers extend from the hinge to the largest value no further than 1.5 times the interquartile range from the hinge. ‘****’ indicates p<0.0001.

This study reveals the potential for bacterial community transplant experiments in freshwater snails. Although such controlled laboratory experiments will undoubtedly provide vital insights into the tripartite interaction between snails, bacteria and parasitic flatworms, extrapolation of patterns and protocols to field settings might prove complicated [5]. However, correlative field-based studies could fill these caveats and, likewise, transplant experiments could help discern correlation from causation in a controlled setting [14]. Correlative studies could help in identifying bacterial strains involved in snail resistance. The potential of these strains could then be further examined in transplant experiments in controlled lab environments and mesocosm studies prior to manipulations in the field. We advocate in doing these contained experiments prior to unconfined biological manipulations in the field as the outcome can be hard to predict. Nevertheless, an improved understanding of the tripartite interaction between hosts, bacteria and pathogens will prove paramount to fully grasp disease epidemiology and field-based correlative studies might help guide experimental efforts.

## Acknowledgements

Our major gratitude goes out to Benjamin Gourbal and the ‘Université de Perpignan Via Domitia’ for providing the lab-reared *Biomphalaria glabrata*. Without these samples our research would not have been possible. We also want to thank Eve Toulza and Bart Lievens for their constructive feedback on the study and methodology design during the PhD committee meetings. Furthermore, our gratitude goes out to Shira Houwenhuyse, Manon Coone, Cyril Hammoud, Amruta Rajarajan and Martijn Callens for providing the foundations of our pipeline. Finally, we want to thank Willem Stock for his critical remarks on our dataset and all who was involved in maintaining the snail culture. The computational resources and services used in this work were provided by the VSC (Flemish Supercomputer Center), funded by the Research Foundation - Flanders (FWO) and the Flemish Government.

## Author contributions

RS, IS and ED conceived the study, collected the samples, and wrote the manuscript. RS and IV conducted the experiments and prepared the sequencing library. RS performed the data analysis and prepared the figures. RS, TH and ED interpreted the data. All authors revised and commented on the manuscript. All authors have read the final version of the manuscript and approved of its content.

## Data Availability Statement

The metabarcoding dataset generated and analysed during the current study will be made available in the NCBI Sequence Read Archive (SRA) repository.

The technique used to obtain microbiome-disturbed snails and rscripts to analyse the dataset will be made available on the Dryad repository.

## References

1. Koch H, Schmid-Hempel P. Gut microbiota instead of host genotype drive the specificity in the interaction of a natural host-parasite system. Ecol Lett 2012; 15: 1095–1103.

2. Macke E, Callens M, De Meester L, Decaestecker E. Host-genotype dependent gut microbiota drives zooplankton tolerance to toxic cyanobacteria. Nat Commun 2017; 8: 1608.

3. Moreira LA, Iturbe-Ormaetxe I, Jeffery JA, Lu G, Pyke AT, Hedges LM, et al. A Wolbachia Symbiont in Aedes aegypti Limits Infection with Dengue, Chikungunya, and Plasmodium. Cell 2009; 139: 1268–1278.

4. Ridaura VK, Faith JJ, Rey FE, Cheng J, Duncan AE, Kau AL, et al. Gut Microbiota from Twins Discordant for Obesity Modulate Metabolism in Mice. Science (80-) 2013; 341: 1241214.

5. Greyson-Gaito CJ, Bartley TJ, Cottenie K, Jarvis WMC, Newman AEM, Stothart MR. Into the wild: microbiome transplant studies need broader ecological reality. Proc R Soc B Biol Sci 2020; 287: 20192834.

6. Ford SA, King KC. Harnessing the Power of Defensive Microbes: Evolutionary Implications in Nature and Disease Control. PLoS Pathog 2016; 12: 1–12.

7. Flandroy L, Poutahidis T, Berg G, Clarke G, Dao M-C, Decaestecker E, et al. The impact of human activities and lifestyles on the interlinked microbiota and health of humans and of ecosystems. Sci Total Environ 2018; 627: 1018–1038.

8. Brinker P, Fontaine MC, Beukeboom LW, Falcao Salles J. Host, Symbionts, and the Microbiome: The Missing Tripartite Interaction. Trends Microbiol 2019; 27: 480–488.

9. Hoffmann AA, Montgomery BL, Popovici J, Iturbe-Ormaetxe I, Johnson PH, Muzzi F, et al. Successful establishment of Wolbachia in Aedes populations to suppress dengue transmission. Nature 2011; 476: 454–457.

10. Pinto SB, Riback TIS, Sylvestre G, Costa G, Peixoto J, Dias FBS, et al. Effectiveness of Wolbachia-infected mosquito deployments in reducing the incidence of dengue and other Aedes-borne diseases in Niterói, Brazil: A quasi-experimental study. PLoS Negl Trop Dis 2021; 15: e0009556.

11. Huot C, Clerissi C, Gourbal B, Galinier R, Duval D, Toulza E. Schistosomiasis Vector Snails and Their Microbiota Display a Phylosymbiosis Pattern. Front Microbiol 2020; 10: 1–10.

12. Le Clec’h W, Nordmeyer S, Anderson TJC, Chevalier FD. Snails, microbiomes, and schistosomes: a three-way interaction? Trends Parasitol 2022; 38: 353–355.

13. Portet A, Toulza E, Lokmer A, Huot C, Duval D, Galinier R, et al. Experimental infection of the Biomphalaria glabrata vector snail by Schistosoma mansoni parasites drives snail microbiota dysbiosis. Microorganisms 2021; 9.

14. Stock W, Callens M, Houwenhuyse S, Schols R, Goel N, Coone M, et al. Human impact on symbioses between aquatic organisms and microbes. Aquat Microb Ecol 2021; 87: 113–138.

15. Sokolow SH, Wood CL, Jones IJ, Lafferty KD, Kuris AM, Hsieh MH, et al. To Reduce the Global Burden of Human Schistosomiasis, Use ‘Old Fashioned’ Snail Control. Trends Parasitol 2018., 34

16. Sène M, Southgate VR, Vercruysse J. Bulinus truncatus, intermediate host of Schistosoma haematobium in the Senegal River Basin (SRB). Bull la Soc Pathol Exot 2004; 97: 29–32.

17. Chernin E. A Method of Securing Bacteriologically Sterile Snails (Australorbis glabratus). Proc Soc Exp Biol Med 1957; 96: 204–210.

18. Chernin E. Infection of Australorbis glabratus with Schistosoma mansoni under Bacteriologically Sterile Conditions. Proc Soc Exp Biol Med 1960; 105: 292–296.

19. Reitmeier S, Hitch TCA, Treichel N, Fikas N, Hausmann B, Ramer-Tait AE, et al. Handling of spurious sequences affects the outcome of high-throughput 16S rRNA gene amplicon profiling. ISME Commun 2021; 1: 31.

20. Zaneveld JR, McMinds R, Vega Thurber R. Stress and stability: applying the Anna Karenina principle to animal microbiomes. Nat Microbiol 2017; 2: 17121.

