## Supplementary information for "Host-microbiome transplants of the schistosome snail host *Biomphalaria glabrata* reflect species-specific associations"

**Materials and Methods**

**Cultivation of *Biomphalaria glabrata***

*Biomphalaria glabrata* (Brazil strain) individuals were kept in 2 l and 5 l aquaria, depending on the cohort size, on a 12/12 day/night cycle at 25°C (clear white light was generated by OSRAM FH 28W/840 HE). Each aquarium had a constant air supply provided through aeration stones. Maternal lines started from one egg clutch and were kept in separate aquaria for 18 months. Fresh biological salad was provided three times per week at libitum. Aquaria were cleaned on a weekly basis with soap and mechanical rubbing, followed by an industrial washing machine program at 55 °C and drying in the oven. Aeration stones were cleaned by boiling in water for 10 min, overnight incubation in diluted bleach (1:3) and, finally, dried by connecting to the air line.

**Sample collection**

Microbiome-disturbed *Biomphalaria glabrata* individuals were exposed to three donor microbiomes across an increasing taxonomic distance. The same species (*B. glabrata*) but different maternal line, the same Family (Planorbidae) *Planorbarius corneus* (collected from an artificial pond with GPS-coordinates: 50.999559, 4.955650), the same Superorder (Hygrophila) *Lymnaea stagnalis* (collected from an artificial pond with GPS-coordinates: 50.915337, 4.686979).

**Transplant experiment**

At the start of each day, egg clutches of each maternal line were collected and assessed for developmental status under a stereo microscope (Olympus SZX10). Late-development clutches were selected, whereby, surrounding eggs were sliced open and debris was removed from around the egg of interest. Egg clusters were dissected for the three maternal lines. Minimally 32 eggs were needed per day, but ±38 were dissected to ensure that sufficient specimens were available during the sterilization experiment. The eggs were kept separately per maternal line in autoclaved tap water (25 °C) in 6-well plates to transport to the sterile biosafety cabinet. Dechlorinated tap water was autoclaved per litre, and is further referred to as “autoclaved tap water”. This water was aliquoted in tubes of 50 ml and was stored next to the snail aquaria at 25 °C. The UV light was turned on 15 min before turning on the circulation in the biosafety cabinet. The working surface and required utensils were disinfected with 70 % ethanol. Bleach was diluted with autoclaved tap water to obtain a working solution of 0.2 % Sodium hypochlorite or 0.042 % active chlorine. Snail eggs would not fit through normal 200 µl tips. Therefore, 200 µl tips were trimmed with sterilized scissors to have a larger opening. All plates required for that day were filled according to **Fig. S1**. Briefly, the second column of the 24-well plates were filled with 1 ml of the 0.2 % bleach solution. All other wells were filled with 1ml of autoclaved tap water. Each first-column well received one dissected egg. Subsequently, eggs were exposed to 0.2 % bleach for 15 min. The timer started when the first egg entered the bleach solution. The following eggs for that plate were added at 30 s intervals. Once all four eggs were in the bleach solution, we would swirl the plate gently for 10 s. The same procedure was started for the second plate one minute after finishing the first plate. This process was repeated for a total of six plates. Next, the first egg, now exposed to bleach for 15 min, was placed in the next well containing autoclaved tap water. We pipetted up and down for 10 s in each well before transferring the egg to the next well to remove any remaining bleach. The first egg was stored in the one-to-last well and we started the same procedure for the second egg. This was repeated for all four eggs of that plate. Now all eggs were placed it in their final wells for hatching. This treatment was repeated for all six plates and the lid-tray interface was closed with parafilm.

To test if 1) treated snails would acquire a given donor microbiome, 2) what the optimal timing for exposure would be and 3) how the phylogenetic relationship between the recipient and donor host species affects the transplant outcome, we designed the following experimental setup (**Fig. 1** main manuscript). Snails were sterilized three days, two days, one day prior to and on the same day as receiving a donor inoculum. Donor inocula from *B. glabrata* and *P. corneus* were obtained by separately crushing three individual snails with a sterile pestle and topping the volume with autoclaved tap water until 1 ml. From each tube, 300 µl was pipetted in a 2 ml tube per donor type topped till 1.4 ml with autoclaved tap water. The same protocol was used for *L. stagnalis*, however, here only two snails were available. From the donor inocula 20 µl was each time added to the desired well. This was done for three separate maternal lines of lab-reared *B. glabrata* each time for four technical replicates (four individual snails were exposed to the same conditions). Each factorial combination (time (n=4)*maternal line (n=3)) received a donor inoculum from, either, a fourth maternal line of lab-reared *B. glabrata*, a wild-caught *P. corneus* and a wild-caught *L. stagnalis*. Additionally, for each factorial combination of time and maternal line a positive (dissected without bleach exposure) and a negative (dissected with bleach exposure but no donor inoculum) control were included. Survival status of the egg/juvenile was noted at the day of the donor inoculum exposure, three days post donor inoculum exposure and four days post donor inoculum exposure when the specimens were sacrificed and the experiment ended. Snails were pooled by up to three specimens per factorial combination (time (n=4) * maternal line (n=3) * donor microbiome (n=3) *survival status (n=3)) in order to obtain sufficient DNA for subsequent analyses. All negative controls and the positive controls of day 0 and day three were tested for the presence of bacteria in the growth assay in combination with microscopic investigation and qPCR. These and all other samples were included for 16S metabarcoding.

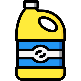

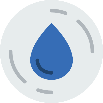

0.2 % bleach solution

Autoclaved tap water

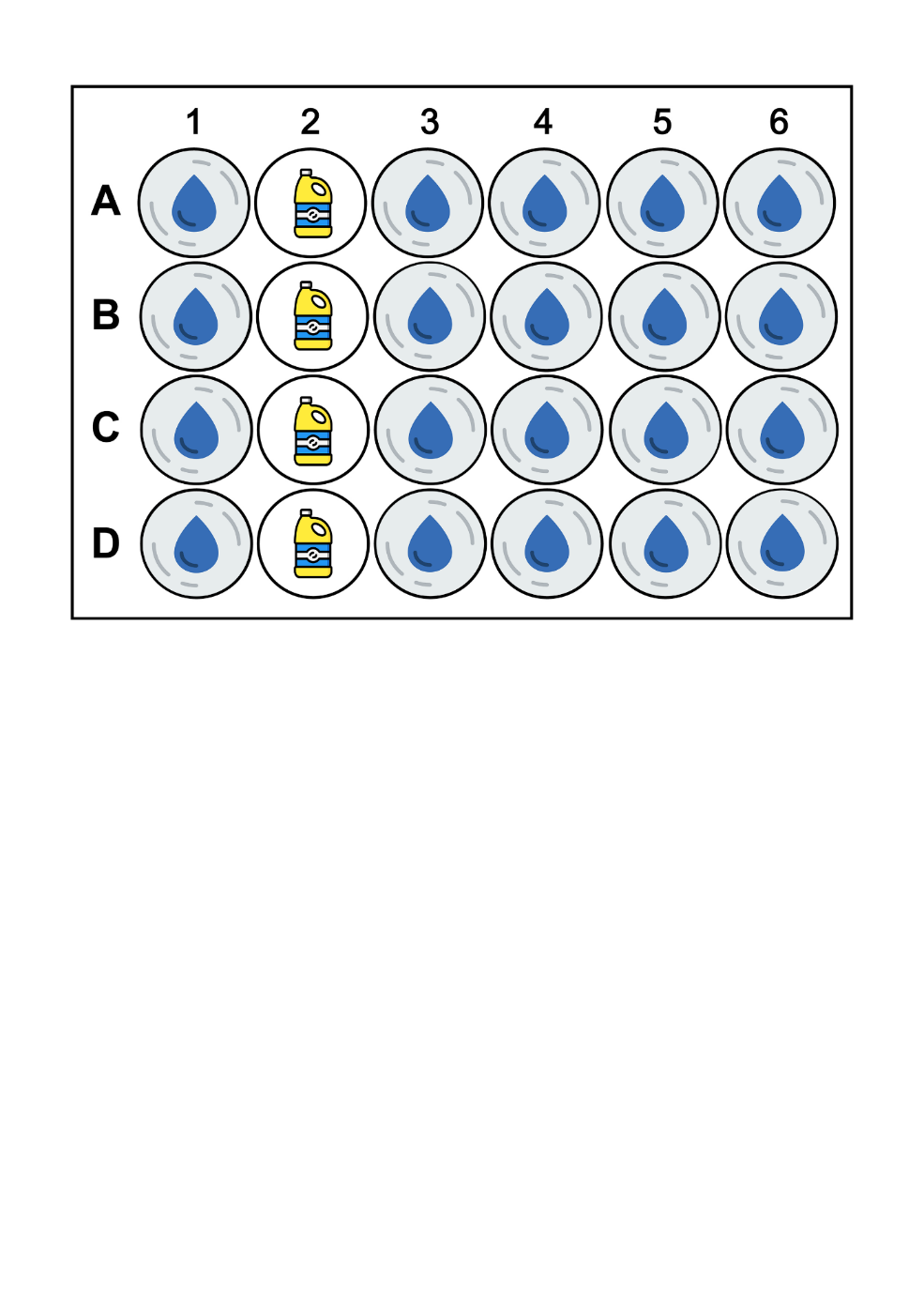

**Fig. S1**: **The 24-well cell culture plate setup used during the sterilization protocol**. The wells of the first column receive one egg each, prior to bleach exposure. The egg from A1 is transferred to the 0.2% bleach containing well in A2 and remains there for 15 min. Before transferring the next egg (from B1 to B2), we waited 30 s. After the 15 min of bleach exposure, eggs were washed in autoclaved tap water in column three, four and five before being moved to their incubation well in column six.

**DNA extraction**

Snails were crushed with a sterile pestle in 100 µl of autoclaved tap water by up to three specimens per factorial combination in order to obtain sufficient DNA for subsequent analyses. The complete homogenate was used for recipients of donor microbiomes. For all negative controls and the positive controls at day zero and three, 90 µl of this homogenate was used for DNA extraction (5 µl was used for microscopic investigation and 5 µl for the growth assay). DNA of all samples was extracted using the E.Z.N.A.® Mollusc DNA Kit (OMEGA Bio-tek, Norcross, GA, USA) according to the manufacturer’s protocol and eluted through two elution steps of 50 μL, totalling to 100 μL of DNA extract. DNA concentrations were determined using the Qubit dsDNA high-sensitivity assay (Thermo Fisher Scientific) for all DNA extracts. 3 ng of template DNA (max. 10 µl) was used for qPCR. This indicated that DNA concentrations were still low for some samples (0.03 ng/ µl). Therefore, the DNA for all samples was concentrated with the Eppendorf™ Concentrator Plus for 30 min at 30 °C, halving the total volume. Subsequently, Qubit measurements were repeated on all samples. This protocol increased the total amount of DNA per PCR reaction from 0.3 ng to 0.6 ng.

**Validating axenicity**

For the growth assay, 5 µl of the homogenate was added to 4 ml of R2A medium. The R2A medium was selected as it is a medium capable of growing a broad range of bacteria (Aditi et al., in prep.). The R2A medium consisted of 0.5 g/l Bacto^TM^ Proteose peptone (Gibco), 0.5 g/l Casamino acids (Gibco), 0.5 g/l Yeast extract (OXOID), 0.5 g/l Dextrose [D-(+)-Glucose] C6H1206 – M 180, 16 g/mol (Carl Roth), 0.5 g/l Soluble starch (Merck), 0.3 g/l Dipotassium hydrogen phosphate (K2HPO4) (Merck), 0.024 g/l Magnesium sulphate (MgSO4.7H20) (Duchefa), 0.3 g/l Sodium pyruvate 100 mM (100x) (Gibco) and topped till one litre with distilled water. The resulting medium was autoclaved and stored at room temperature. The sterile 14 mL tube (Greiner Bio-one) was vortexed briefly and incubated with constant mechanical mixing at 25 °C for 72 h. Prior to OD measurements at 600 nm, the R2A medium was pipetted up and down several times to ensure adequate mixing and homogeneous OD values per sample. Measurements were done on 1 mL of medium in the Genesys 10S UV-Vis Spectrophotometer (Thermo Fisher Scientific) using the Macro-cuvette, 1 mL (Greiner Bio-one). R2A medium incubated with 5 µl of autoclaved tap water was used as a negative to obtain 0 OD values.

For the microscopic investigation for the presence of bacteria, the LIVE/DEAD® *Bac*Light^TM^ Bacterial Viability Kit L13152 (Thermo Fisher Scientific) was used. Each dye was dissolved in 2.5 ml of autoclaved tap water and 10 µl of a (2/3 Propidium Iodide and 1/3 SYTO9) mix to 5 µl of sample. Then it was incubated in the dark for 15 min. Subsequently, 6 µl of the sample was placed under a cover glass and the slide was visually inspected under a fluorescence microscope (Olympus BX51, U-MWB2 Blue fluorescence filter, metal halide lamp). Twenty fields of view were examined for bacterial presence at random, unless a piece of tissue/shell/matrix was seen in the field of view, then this received special attention as bacteria were notably more likely to grow un such substrates, thus increasing our chances of finding bacteria in supposedly microbiome-disturbed samples.

qPCR reactions were run on the LightCycler® 480 (Roche). Each reaction consisted of 10 µl of LightCycler® 480 SYBR Green I Master (Life Science), 1 µl 16S-Fw (5’-AGACACGGTCCAGACTCCTAC-3’) at 10µM, 1 µl 16S-Rv (5’-CTTGCACCCTCCGTATTACCG-3’) at 10µM, 3ng of template DNA (n µl) and 8-n µl of sterile Milli-Q water to a final reaction volume of 20 µl [1]. The temperature cycle was as follows, 5min of initial denaturation at 95°C, followed by 45 cycles of 10 sec at 95°C, 20 sec at 60°C and 5 s at 72°C [1]. Each sample was done in triplicate. Standards were obtained by extracting DNA from a pure *E. coli* culture, measuring the DNA concentration and comparing this weight to the weight of a single *E. coli* genome [1]. Standards of the desired concentration were mixed with 4 µl of salmon sperm in a total volume of 5 µl. The melting and amplification curves were calculated through the LightCycler® 480 software (v. 1.5.1) and exported as reports to pdf and excel. The excel dataset was then further analysed in R .

**Mock communities**

In order to optimize and quantify any biases during our pipeline, two types of mock communities were included in the 16S metabarcoding run. First, the 10 Strain Even Mix Genomic Material MSA-1000™ (ATCC) is a DNA-based mock community that allowed us to exclude any potential extraction bias. It consists of 10% *Bacillus pacificus* (ATCC 10987), 10% *Bifidobacterium adolescentis* (ATCC 15703), 10% *Clostridium beijerinckii* (ATCC 35702), 10% *Deinococcus radiodurans* (ATCC BAA-816), 10% *Enterococcus faecalis* (ATCC 47077), 10% *Escherichia coli* (ATCC 700926), 10% *Lactobacillus gasseri* (ATCC 33323), 10% *Cereibacter sphaeroides* (ATCC 17029), 10% *Staphylococcus epidermidis* (ATCC 12228) and 10% *Streptococcus mutans* (ATCC 700610). Second, the 10 Strain Even Mix cell Material MSA-1000™ (ATCC) is a cell-based mock community that allowed us to assess any potential extraction bias by comparing the results to the DNA-based mock community (2x107 whole cells/vial ± 1 log). It consists of freeze dried cell material of the same 10 strains and ratios as the MSA-1000™ mock community. The mock communities also allow us to determine the suitable threshold below which spurious sequences should be removed as discussed by Reitmeier et al. [2].

**16S metabarcoding**

The PCR protocol targeting the V3 and V4 hypervariable regions of the 16S rRNA gene [3] was adapted from [4]. PCR reactions were conducted in triplicate for all samples in a 25 µl total volume. 12.5 µl of PCRBIO HS VeriFi^TM^ Mix (PCR Biosystems^TM^), 0.75 µl of the primer 341F (5’-CCTACGGGNGGCWGCAG-3’) at 20 µM, 0.75 µl of the primer 805R (5’-GACTACHVGGGTATCTAATCC-3’) at 20 µM and a variable volume of DNA (0.6 ng/DNA concentration of sample) and water (11 µl – the volume of DNA) to complete the volume to 25 µl and obtain a total DNA input of 0.6 ng. An initial denaturation at 98 °C was done for 5 min, followed by 25 cycles of denaturation at 98 °C for 45 s, annealing at 60 °C for 15 s and extension at 72 °C for 10 s and, finally, an elongation step at 72 °C for 7 min. The resulting triplicate PCR products were pooled per sample and subsequently purified with magnetic beads (CleanNGS, GC Biotech) following the manufacturer’s protocol. The purified PCR products were sent to the Genomics core at UZ Leuven for sequencing (Illumina Miseq v3). First a quality control was performed through a fragment analyzer, and the DNA concentration was measured through Qubit. Then a second PCR reaction attached the Illumina adapters and indexes to the DNA fragments through the Genomics core tails. This PCR was done in a total volume of 20 µl and consisted of 9 µl PCR1 product (concentration 0.5-5 ng/ µl), 0.5 µl of the forward P7 primer (5 µM), 0.5 µl of the Reverse P5 primer (5 µM) and 10 µl of Phusion high fidelity PCR master mix (Thermo Fisher Scientific). PCR conditions were the following. First an initial denaturation step at 94 °C for 30 s, 15 cycles of a denaturation step at 94 °C for 10 s, an annealing step at 51 °C for 30 s and an elongation step at 72 °C for 30 s and, finally, an elongation step at 72 °C for 1 min. After this PCR a second quality control was done with the fragment analyzer to make sure the adapters have joined properly (i.e. fragments increased by +-60bp). Qubit analysis was performed once more and a final equimolar library was created. Finally, 14 pM was loaded with 37 % PhiX spike-in on the Illumina MiSeq v3 sequencer (2 x 300 bp with the 600-cycle kit).

**Data analyses**

The QIIME2 pipeline v2022.2 [5] was used to process the raw sequence data based on [6]. First, the paired-end demultiplexed sequence reads were imported and their quality scores were assessed through MultiQC [7]. The run resulted in 4 708 922 raw sequences (min 22 608, max 142 259), 123 622 allocated to mock communities. Using the denoise function of DADA2 within QIIME2, both forward and reverse reads were trimmed for 15 bp and truncated at 280 bp for the forward read and 240 bp for the reverse read, while demanding a 30 bp overlap between both reads. Furthermore, the paired-end sequences were filtered, denoised and dereplicated, chimeras were filtered and an amplicon sequence variants (ASVs) table was created using this denoise function. As a result, on average 59.4% of the reads were maintained per sample (sd= 9.7%). A total of 3 161 ASVs across all samples was obtained with an average length of 425.82 bp (sd= 12.27). The Silva 138 SSU Ref NR 99 database [8] was used to assign taxonomy to the ASVs. It contained the extracted sequences of the V3-V4 region using the forward primer (5’-CCTACGGGNGGCWGCAG-3’), reverse primer (5’-GACTACHVGGGTATCTAATCC-3’) and truncation length (465 bp). The R pipeline of [9] was modified for this study and ran with R (v. 4.2.2, 64-bit) using RStudio (v. 2022.12.0) on a Windows machine (Windows 11). Briefly, the resulting feature table, taxonomy, metadata and phylogenetic tree were imported in R through the phyloseq package (v. 1.42.0, [10]). The R package Decontam (version: 1.18.0; [11]) was used to look for contaminating DNA using the prevalence method and a 0.1 threshold. The PCR negative and extraction negative were used as negative samples to detect contamination. The mock community dataset did not have any contamination ASVs from the total 158 ASVs. The transplant experiment had 64 contaminating ASVs out of the total 3 156 ASVs removed from the dataset. Cleaning and filtering of the dataset was performed in R (see suppl. file for details). The Vegan package (v. 2.6-4, [12]) was used to construct a rarefaction species richness curve [13]. Samples were rarefied to the sampling depth of the sample with the fewest reads (i.e. 15 321) using the “rarefy_even_depth” function of the phyloseq package (rngseed = 711). Consequently, removing 208 ASVs from the dataset. Subsequently, low abundance taxa (< 0.5%, 2 046 ASVs; see below) were removed through the “prune_taxa” function of the phyloseq package. Alpha diversity measures included the Shannon (phyloseq package, function “estimate_richness”) and the Faith's phylogenetic diversity (picante package, v. 1.8.2 [14]; function “pd”). Beta diversity measures included the Bray Curtis and Jaccard index, visualized through NMDS (phyloseq package, function “ordinate”) and RDA (vegan package, function “capscale”). The micrUBIfuns (v. 0.1.0.) function was modified to create **Fig. 2b** (main manuscript) and study the within group beta diversity for several diversity measures. Core microbiome analyses were conducted with the microbiome package (v. 1.20.0, [15]) using different detection (0.01 – 0.001) and prevalence (0.85 – 0.95, to allow for each subset a maximum of one sample without the ASV) values. Kaplan-Meier survival curves were created (survival package, v. 3.4-0; function “survfit”) and significance tested (survival package, function “survdiff”). Finally, to assess a suitable threshold below which spurious sequences should be removed, a maximum likelihood (ML) tree was calculated through the IQ-tree web servers (http://iqtree.cibiv.univie.ac.at), and nodal support was assessed through 1 000 bootstrap replicates [16]. The resulting tree was midpoint rooted in Figtree (version 1.4.4) and tip labels were edited. The resulting tree was exported in PDF format to Adobe Acrobat Pro DC (version 2022.003.20310), in which we adjusted the position of ill-placed nodal support values and italicized species names.

**Supplementary results**

OD measurements indicate that six out of 12 samples were germ-free (**Table S1**). The staining method obtained similar results but results were less conclusive. qPCR indicated that Cp values of five samples were not significantly different from Milli-Q water and thus can be considered germ-free. Notably, the growth and microscopic assay detected one additional sample as germ-free while qPCR indicated high bacterial contamination in this sample (Cp value lower than the 10^4 standard).

**Table S1**: **Results from the axenicity validation**. Optical density measurements at 600 nm (OD), staining method (Syto 9) and qPCR (Cp) were done on the negative and positive samples. The question mark indicates samples where the interpretation was inconclusive. The ‘*’ indicates samples that had Cp values not-significantly different from qPCR negatives.

| Sample | OD | Syto 9 | qPCR (Cp+-sd) |
| --- | --- | --- | --- |
| 0A7- | 0.178 | bacteria | 30.08 +- 0.27 |
| 0A8- | 0.028 | bacteria but most look dead | 30.24 +- 0.01 |
| 0A2- | 0 | no bacteria | 32.58 +- 0.41* |
| 24A7- | 0.359 | some bacteria | 26.43 +- 0.14 |
| 24A8- | 0 | no bacteria (?) | 33.03 +- 0.22* |
| 24A2- | 0.428 | some bacteria | 28.04 +- 0.05 |
| 48A7- | 0 | no bacteria (?) | 21.31 +- 0.17 |
| 48A8- | 0 | no bacteria (?) | 34.06 +- 0.60* |
| 48A2- | 0 | no bacteria | 34.28 +- 0.32* |
| 72A7- | 0.417 | bacteria | 19.99 +- 0.21 |
| 72A8- | 0 | no bacteria (?) | 32.71 +- 0.11* |
| 72A2- | 0.005 | bacteria | 30.52 +- 0.18 |
| 0A7+ | 0.52 | bacteria | 25.99 +- 0.09 |
| 0A8+ | 0.507 | bacteria | 24.08 +- 0.02 |
| 0A2+ | 0.406 | bacteria | 24.10 +- 0.14 |
| 72A7+ | 0.367 | bacteria | 28.65 +- 0.49 |
| 72A8+ | 0.361 | bacteria | 26.14 +- 0.32 |
| 72A2+ | 0.382 | bacteria | 27.61 +- 0.19 |
| 10^1 | / | / | 30.56 +- 0.12 |
| 10^2 | / | / | 29.99 +- 0.22 |
| 10^3 | / | / | 27.43 +- 0.17 |
| 10^4 | / | / | 23.83 +- 0.09 |
| Milli-Q1 | / | / | 33.89 +- 0.13 |
| Milli-Q2 | / | / | 33.63 +- 0.23 |

There was a significant difference in Cp values between the samples (p<001) (**Fig. S2**). A highly significant effect (p<0.001) was detected for treatment, maternal line and the interaction between treatment and maternal line (**Fig. S3**). No effect of time was detected on the Cp values. Notably, maternal line 7 was associated with the lowest Cp values (highest bacterial load, **Fig. S3**). The effect remained when we corrected the interaction of the maternal line and treatment with the time of dissection.

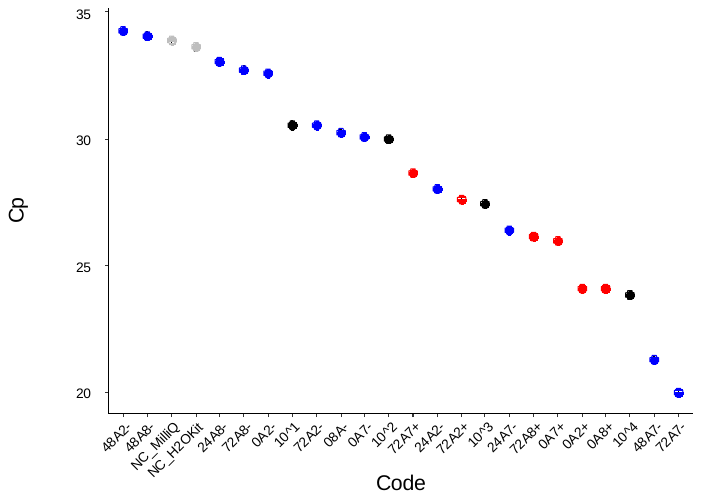

**Fig. S2**: **Cp values for all samples tested in qPCR.** Sample codes were constructed as follows. First, 0, 24, 48, and 72 indicate whether the sample was made microbiome-disturbed at the day of, or, one, two or three days prior to inoculum exposure. Second, A2, A7 and A8 refer to either of the three maternal lines used during this experiment. Finally, ‘+’ or ’-‘ refers to the sample being a positive or a negative control (without bleach exposure or without donor inoculum exposure, respectively). Negative controls, treated samples, positive samples and standards are, respectively, colored as grey, blue, red and black. Standards contained 10^1^, 10^2^, 10^3^ and 10^4^ genomic copies of *E. coli.*

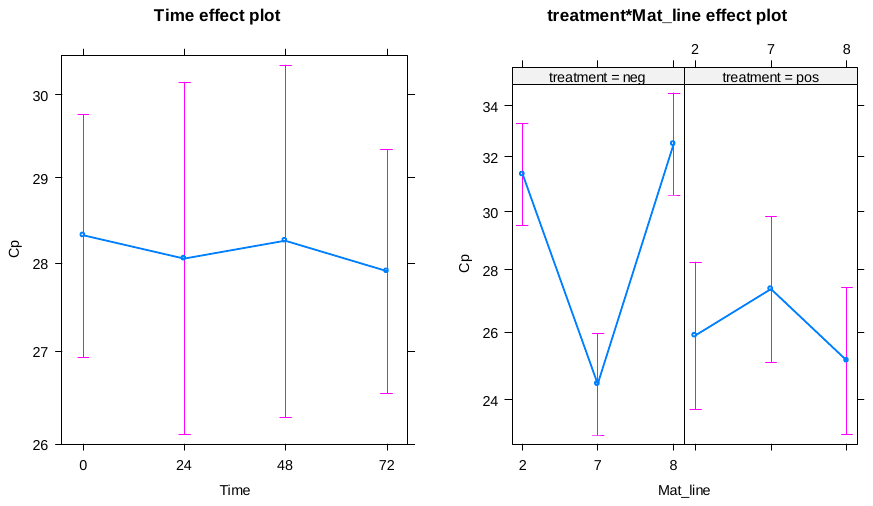

**Fig. 3S: Cp values for the interaction effect between maternal line and the treatment. ‘**Neg’ refers to negative controls, ‘pos’ to positive controls (i.e. without donor inoculum exposure or without bleach exposure, respectively).

**Kaplan-Meier survival curves**

Across time and maternal lines, there was a significant effect of treatment on survival rate of juvenile snails (**Fig. S4**). Moreover, dissected but non-sterilized snails (pos) survived significantly better than any other treatment. Snails that received *P. corneus* donor microbiome (PLA) survived significantly better than snails without (neg) or *L. stagnalis* (LYM) donor microbiomes (p=0.02). A similar yet unsignificant trend is notable for snails that received *B. glabrata* donor microbiome (GLA) (p=0.07). Sterilization at the same day as exposure to a donor microbiome induced high mortality rates (**Fig. S5**). There was a significantly reduced survival rate for GLA, PLA, LYM and neg samples at day 0 (**Fig. S5**).

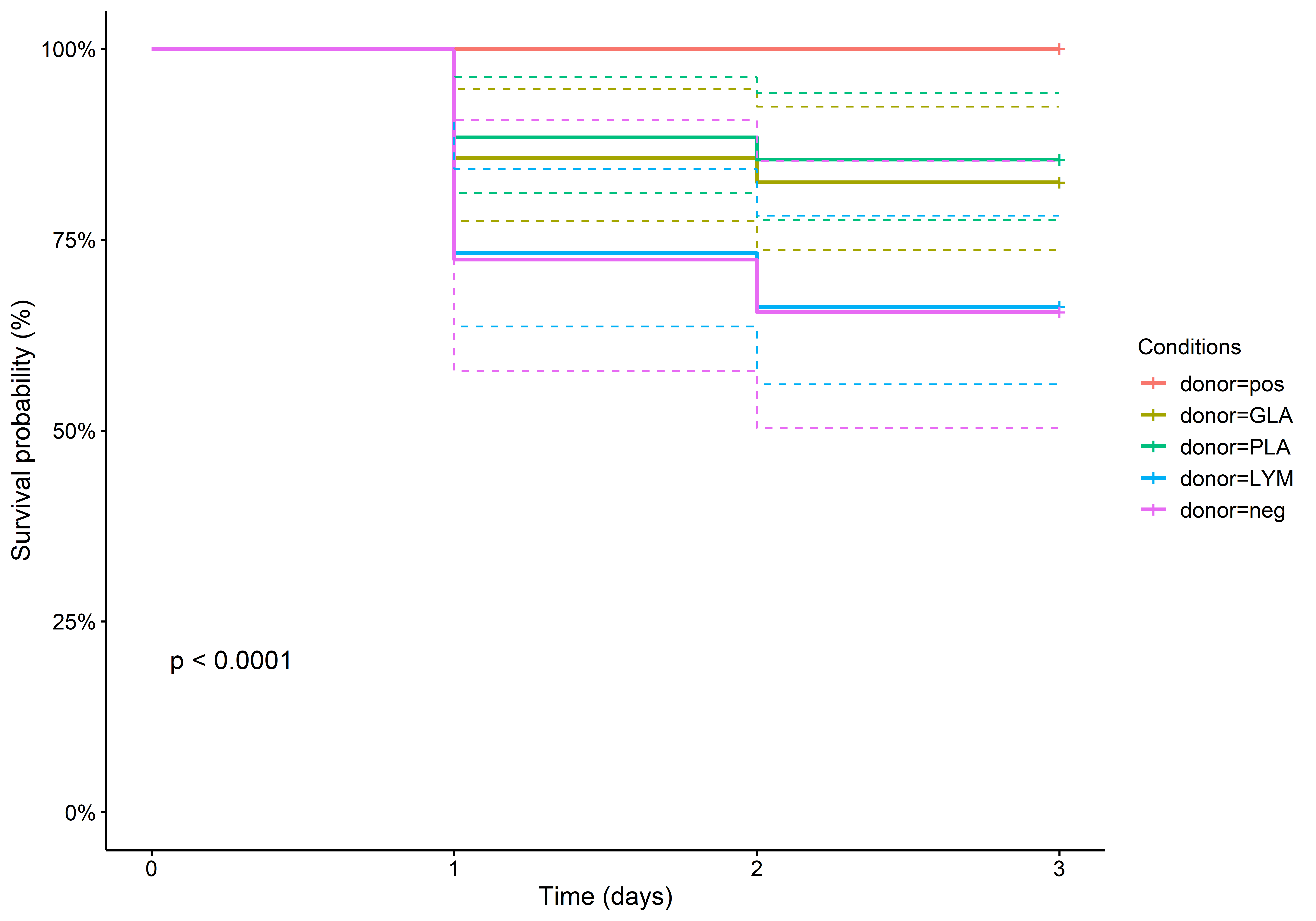

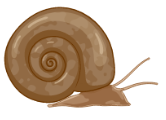

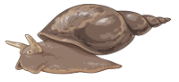

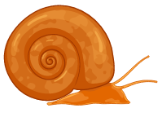

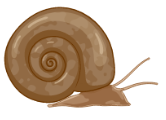

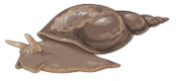

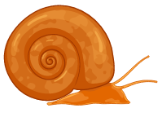

**Fig. S4: Kaplan-Meier survival curve of the complete model (across all days). ‘**Pos’ refers to positive controls (without bleach exposure), ‘GLA’ indicates the samples that received a donor inoculum from *B. glabrata*, ‘PLA’ indicates the samples that received a donor inoculum from *P. corneus*, ‘LYM’ indicates the samples that received a donor inoculum from *L. stagnalis* and ‘neg’ refers to negative controls (without donor inoculum exposure). The difference in survival probability between the different donor treatments was highly significant (p<0.0001).

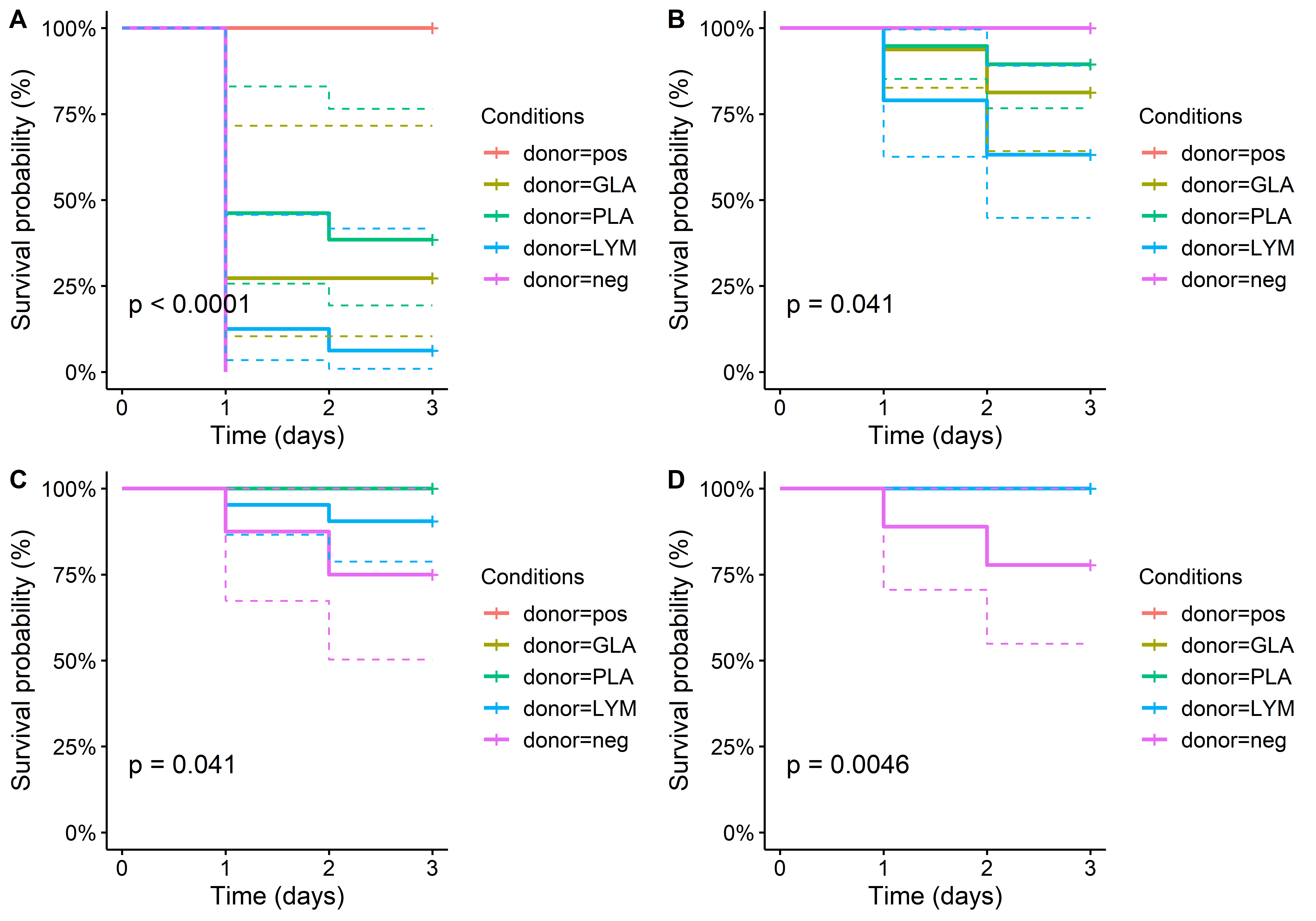

**Fig. S5:** The survival probability of all treatments across the days between sterilization and inoculum exposure. A) Sterilization and inoculum exposure on the same day. B) Sterilization one day before inoculum exposure. C) Sterilization two days before inoculum exposure. D) Sterilization one day before inoculum exposure. ‘Pos’ refers to positive controls (without bleach exposure), ‘GLA’ indicates the samples that received a donor inoculum from *B. glabrata*, ‘PLA’ indicates the samples that received a donor inoculum from *P. corneus*, ‘LYM’ indicates the samples that received a donor inoculum from *L. stagnalis* and ‘neg’ refers to negative controls (without donor inoculum exposure). The difference in survival probability between the different donor treatments was significant for all dissection days.

**16S metabarcoding**

**Mock communities**

We aimed to remove as many spurious sequences as possible through the use of bacterial mock communities according to Reitmeier et al. [2]. These results indicated that a threshold of 0.5 % was most appropriate for the cumulative read abundance of our dataset (**Fig. S6, Table S2**).
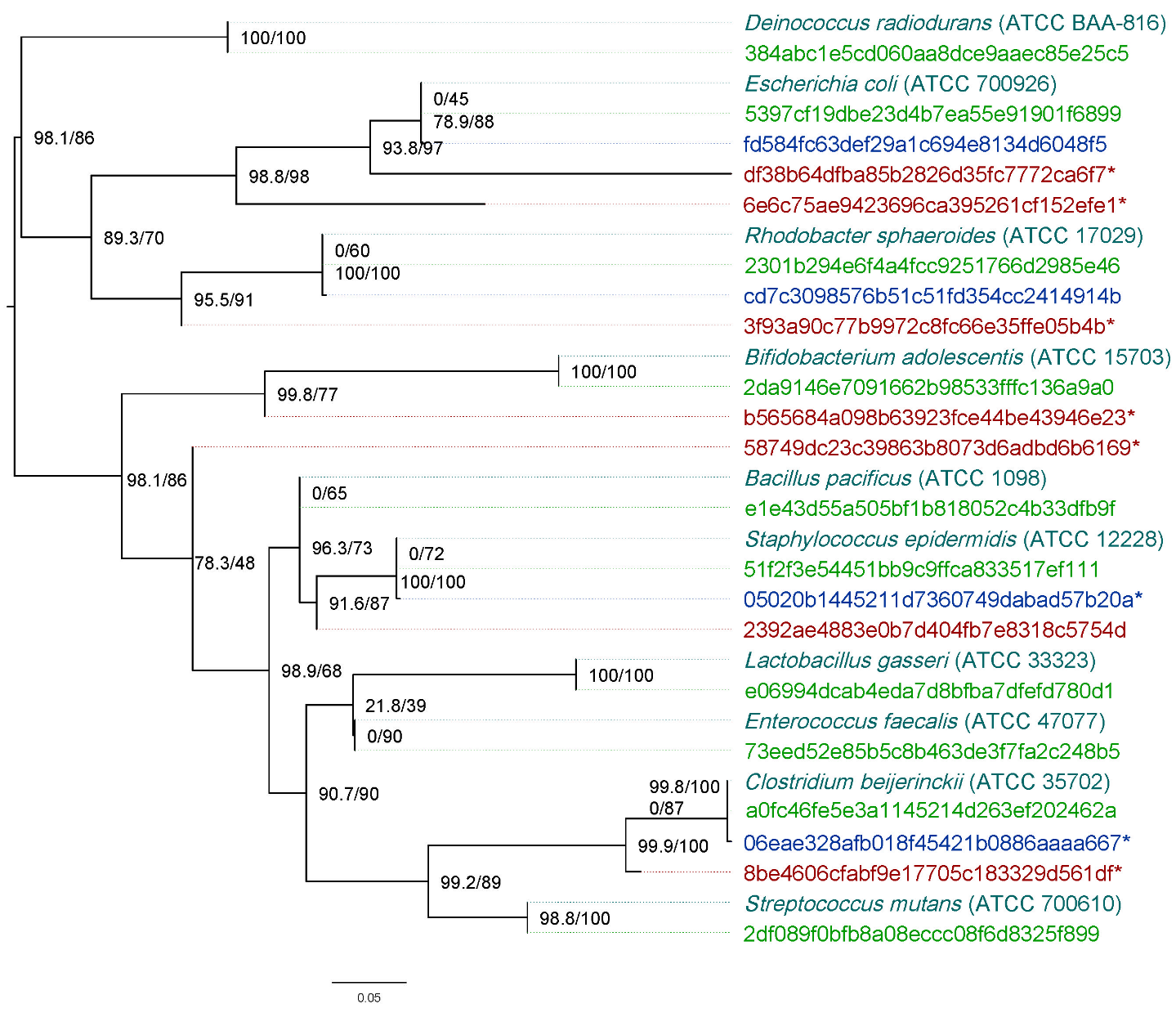

**Fig. S6**: Maximum likelihood midpoint rooted phylogenetic tree using 16S rDNA (453 bp) and using the K3P+ G model (G = 0.46). Nodal support is indicated as bootstrap percentages (1 000 bootstraps, before the ‘/’). Tree branch support by SH-like aLRT with 1 000 replicates (after the ‘/’). Cyan indicates reference sequences from both DNA and cell-based mock communities. Their respective identifier from the ATCC website is provided between parenthesis. Green indicates ASVs perfectly matching the respective reference sequence. Blue indicates at least one bp mismatch with the respective reference sequence. Red indicates ASVs that do not cluster with any reference sequence. ASVs removed after increasing the relative abundance threshold to 0.5 % are indicated by ‘*’.

**Table S2: Cumulative read abundance (%) of each ASV across all mock community samples.** ASVs exceeding the 0.5% threshold are indicated as such in the last column. °indicates a samples that would have ideally been removed from the dataset based on **Fig. S6**. However, increasing the threshold further would risk losing valid data.

| **ASV** | **Cumulative relative read abundance (%) across mock communities** | **Exceeds threshold?** |
| --- | --- | --- |
| b565684a098b63923fce44be43946e23 | 0.028 | No |
| e1e43d55a505bf1b818052c4b33dfb9f | 52.031 | Yes |
| 2392ae4883e0b7d404fb7e8318c5754d | 0.642 | Yes° |
| 73eed52e85b5c8b463de3f7fa2c248b5 | 0.926 | Yes |
| e06994dcab4eda7d8bfba7dfefd780d1 | 3.599 | Yes |
| 2df089f0bfb8a08eccc08f6d8325f899 | 14.714 | Yes |
| 51f2f3e54451bb9c9ffca833517ef111 | 1.147 | Yes |
| 05020b1445211d7360749dabad57b20a | 0.106 | No |
| 58749dc23c39863b8073d6adbd6b6169 | 0.014 | No |
| 8be4606cfabf9e17705c183329d561df | 0.013 | No |
| a0fc46fe5e3a1145214d263ef202462a | 19.361 | Yes |
| 06eae328afb018f45421b0886aaaa667 | 0.305 | No |
| 384abc1e5cd060aa8dce9aaec85e25c5 | 19.938 | Yes |
| 2301b294e6f4a4fcc9251766d2985e46 | 135.290 | Yes |
| cd7c3098576b51c51fd354cc2414914b | 2.689 | Yes |
| 3f93a90c77b9972c8fc66e35ffe05b4b | 0.050 | No |
| 2da9146e7091662b98533fffc136a9a0 | 1.203 | Yes |
| 5397cf19dbe23d4b7ea55e91901f6899 | 244.204 | Yes |
| fd584fc63def29a1c694e8134d6048f5 | 3.707 | Yes |
| df38b64dfba85b2826d35fc7772ca6f7 | 0.013 | No |
| 6e6c75ae9423696ca395261cf152efe1 | 0.019 | No |

**Alpha diversity**

There was a significant difference (p< 0.005) between the treatment types when assessing the complete model (**Fig. S7**). The PCR negative, samples that were sterilized but remained without donor microbiome (neg) and the *L. stagnalis* donor had a significantly lower alpha diversity compared to the other treatments (p<0.01). The Shannon diversity indicated a significant difference between the negative samples and the extraction negatives in contrast to the Faiths PD value (**Fig. S7**). Samples that were inoculated with *B. glabrata* or *P. corneus* were not significantly different from their respective donor inocula. In contrast, samples that were exposed to the *L. stagnalis* donor inoculum were significantly different from the donor. This discrpancy is presumed to be a consequence of a sequencing bias of the *L. stagnalis* donor inoculum as quality checks prior to sequencing indicated failed amplification but the sample was included nevertheless.

**Fig S7: Alpha diversity across the transplant experiment**. **a** Shannon diversity index **b** Faith’s PD per treatment group. GLA indicates the samples that received *B. glabrata* (different maternal line) donor microbiome, PLA indicates the samples that received *P. corneus* donor microbiome, LYM indicates the samples that received *L. stagnalis* donor microbiome.

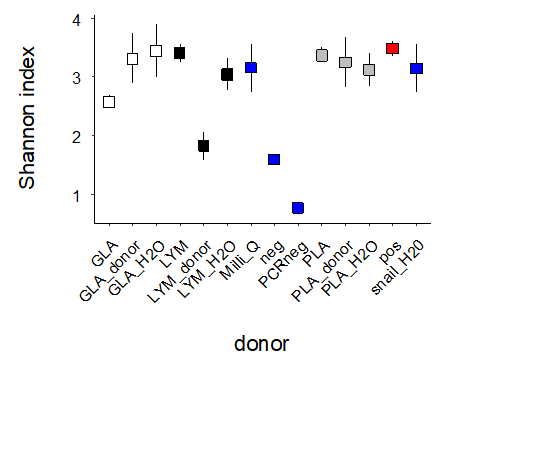

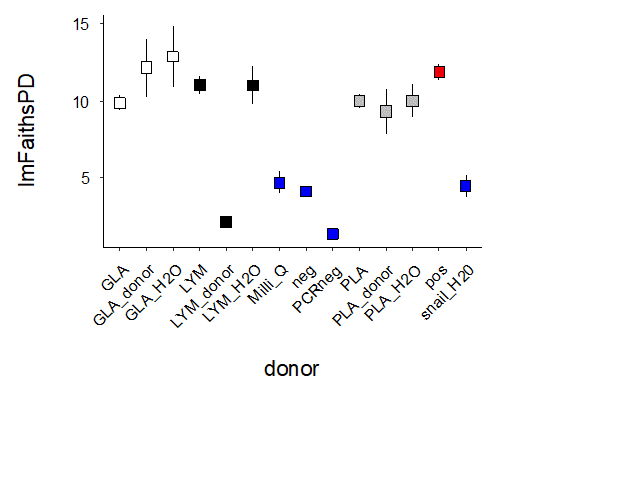

**a**

**b**

**Beta diversity**

Bray-Curtis dissimilarity indicates that all treatments affected the beta diversity of samples differently (**Fig. S8**). The high variation between samples of the ‘neg’ treatment corroborates the high within treatment variation shown in **Fig. 2b** of the main manuscript. The pattern of **Fig. 2a** of the main manuscript remains when considering the Jaccard, Manhattan, unweighted Unifrac or Weighted Unifrac distance measures (**Fig S9**).

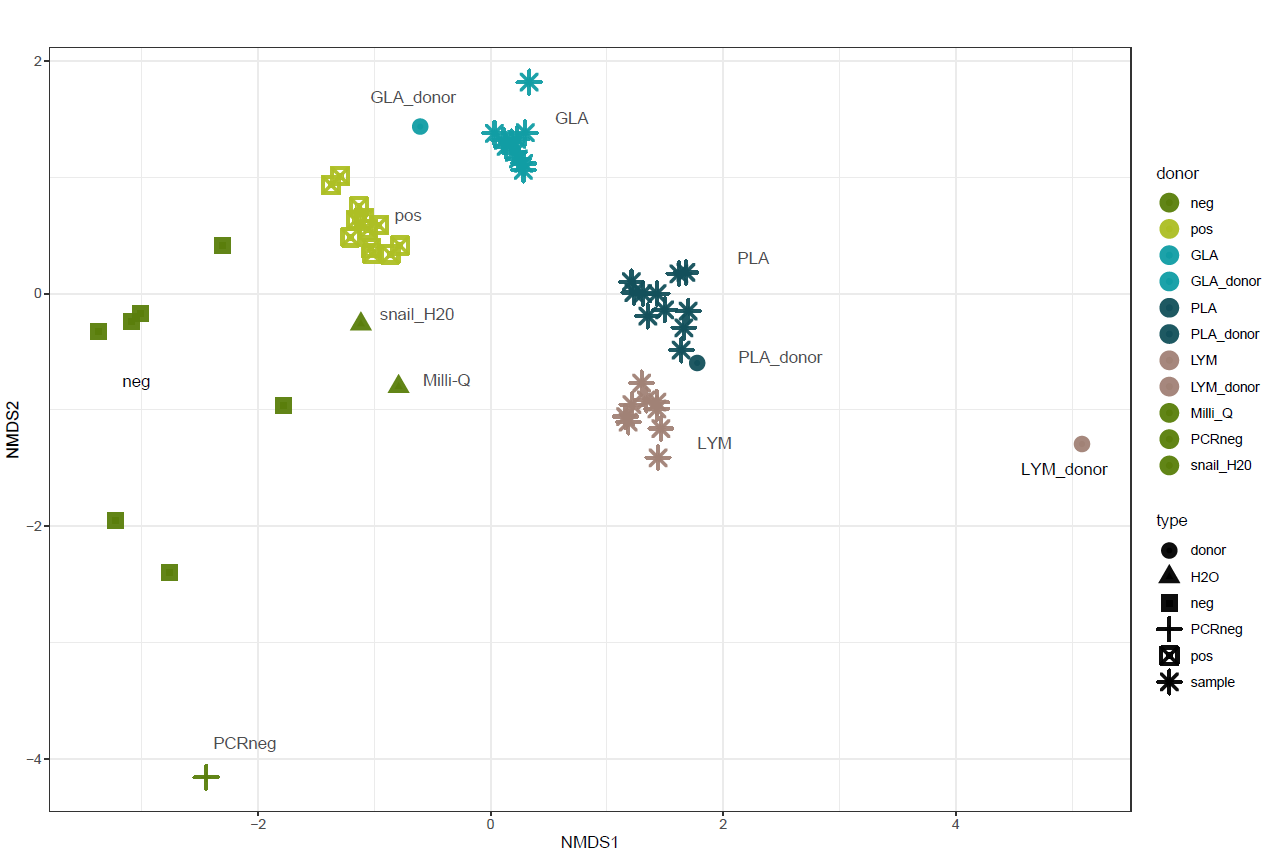

**Fig. S8: NMDS plot showing the Bray-Curtis dissimilarity across all samples**. ‘pos’ refers to positive controls (without bleach exposure). ‘GLA’ indicates the samples that received a donor inoculum from *B. glabrata*. ‘PLA’ indicates the samples that received a donor inoculum from *P. corneus*. ‘LYM’ indicates the samples that received a donor inoculum from *L. stagnalis*. ‘neg’ refers to negative controls (without donor inoculum exposure). ‘Milli-Q’ refers to a Milli-Q extraction using the EZNA mollusc kit used for all DNA extractions. ‘snail_h20’ refers to an extraction done on autoclaved tap water. ‘PCRneg’ is the PCR negative included in sequencing.

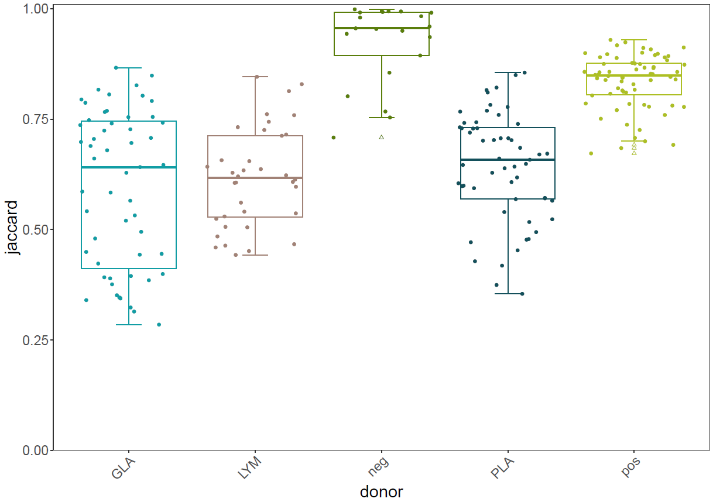

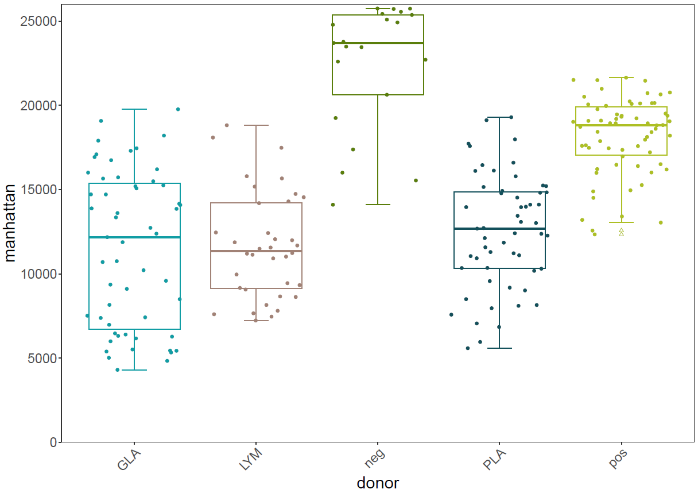

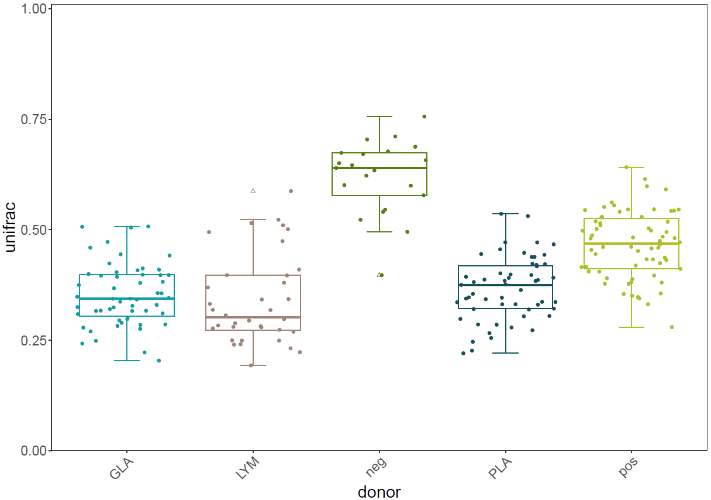

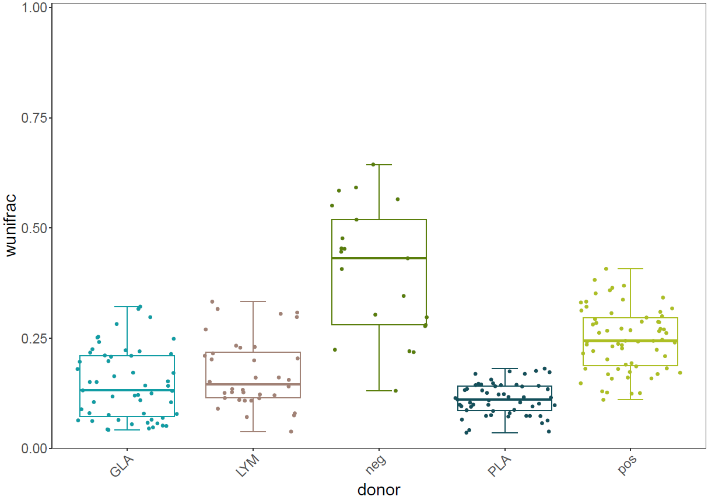

**a**

**b**

**c**

**d**

**Fig S9: The within group beta dissimilarities (n=233)**. **a** Jaccard coefficient. **b** Manhatten distance. **c** unweighted UniFrac distance. **d** weighted UniFrac distance. The abbreviations refer to the following: GLA (*Biomphalaria* *glabrata* recipients), LYM (*Lymnaea* *stagnalis* recipients), neg (microbiome-disturbed specimens), PLA (*Planorbarius* *corneus* recipients) and pos (untreated specimens). Outliers are indicated by triangles. The lower and upper hinges correspond to the first and third quartiles (the 25th and 75th percentiles).

**Core microbiome**

The samples that received *B. glabrata, P. corneus* or *L. stagnalis* donor inocula all shared ASVs belonging to the Family Pseudomonaceae, Comamonadaceae and Flavobacteriaceae when considering a prevalence of minimally 95 % (max. missing in one sample) and a relative read abundance of minimally 1 %. These core members made up 49 % of the reads across all samples (sd= 12 %). ASVs belonging to the Family Pirellulaceae were part of the core microbiome of microbiome-disturbed snails and occurred in all but one sample. This core member had an abundance of 32 % (sd= 28 %) across all negative samples. It also did not occur in the PCR negative, indicating that it is not due to post-experiment contamination.

**References**

1.	Callens M, Watanabe H, Kato Y, Miura J, Decaestecker E. Microbiota inoculum composition affects holobiont assembly and host growth in Daphnia. *Microbiome* 2018; **6**: 56.

2.	Reitmeier S, Hitch TCA, Treichel N, Fikas N, Hausmann B, Ramer-Tait AE, et al. Handling of spurious sequences affects the outcome of high-throughput 16S rRNA gene amplicon profiling. *ISME Commun* 2021; **1**: 31.

3.	Klindworth A, Pruesse E, Schweer T, Peplies J, Quast C, Horn M, et al. Evaluation of general 16S ribosomal RNA gene PCR primers for classical and next-generation sequencing-based diversity studies. *Nucleic Acids Res* 2013; **41**: e1–e1.

4.	Huot C, Clerissi C, Gourbal B, Galinier R, Duval D, Toulza E. Schistosomiasis Vector Snails and Their Microbiota Display a Phylosymbiosis Pattern. *Front Microbiol* 2020; **10**: 1–10.

5.	Bolyen E, Rideout JR, Dillon MR, Bokulich NA, Abnet CC, Al-Ghalith GA, et al. Reproducible, interactive, scalable and extensible microbiome data science using QIIME 2. *Nat Biotechnol* 2019; **37**: 852–857.

6.	Janssens L, Van de Maele M, Delnat V, Theys C, Mukherjee S, De Meester L, et al. Evolution of pesticide tolerance and associated changes in the microbiome in the water flea Daphnia magna. *Ecotoxicol Environ Saf* 2022; **240**: 113697.

7.	Ewels P, Magnusson M, Lundin S, Käller M. MultiQC: summarize analysis results for multiple tools and samples in a single report. *Bioinformatics* 2016; **32**: 3047–3048.

8.	Quast C, Pruesse E, Yilmaz P, Gerken J, Schweer T, Yarza P, et al. The SILVA ribosomal RNA gene database project: improved data processing and web-based tools. *Nucleic Acids Res* 2013; **41**: D590–D596.

9.	Callahan BJ, Sankaran K, Fukuyama JA, McMurdie PJ, Holmes SP. Bioconductor Workflow for Microbiome Data Analysis: from raw reads to community analyses. *F1000Research* 2016; **5**.

10.	McMurdie PJ, Holmes S. phyloseq: An R Package for Reproducible Interactive Analysis and Graphics of Microbiome Census Data. *PLoS One* 2013; **8**: e61217.

11.	Davis NM, Proctor DM, Holmes SP, Relman DA, Callahan BJ. Simple statistical identification and removal of contaminant sequences in marker-gene and metagenomics data. *Microbiome* 2018; **6**: 226.

12.	Oksanen J, Blanchet FG, Kindt R, Legendre P, Minchin PR, O’hara RB, et al. Community ecology package. *R Packag version* 2013; **2**: 321–326.

13.	Heck Jr. KL, van Belle G, Simberloff D. Explicit Calculation of the Rarefaction Diversity Measurement and the Determination of Sufficient Sample Size. *Ecology* 1975; **56**: 1459–1461.

14.	Kembel SW, Cowan PD, Helmus MR, Cornwell WK, Morlon H, Ackerly DD, et al. Picante: R tools for integrating phylogenies and ecology. *Bioinformatics* 2010; **26**: 1463–1464.

15.	Lahti L, Shetty S. microbiome R package. 2012.

16.	Trifinopoulos J, Nguyen L-T, von Haeseler A, Minh BQ. W-IQ-TREE: a fast online phylogenetic tool for maximum likelihood analysis. *Nucleic Acids Res* 2016; **44**: W232–W235.
